# Viral infection enhances vomocytosis of intracellular fungi via Type I interferons

**DOI:** 10.1101/512293

**Authors:** Paula I Seoane, Leanne M. Taylor-Smith, David Stirling, Lucy C. K. Bell, Mahdad Noursadeghi, Dalan Bailey, Robin C. May

## Abstract

*Cryptococcus neoformans* is an opportunistic human pathogen, which causes serious disease in immunocompromised hosts. Infection with this pathogen is particularly relevant in HIV+ patients, where it leads to around 200,000 deaths *per annum*. A key feature of cryptococcal pathogenesis is the ability of the fungus to survive and replicate within the phagosome of macrophages, as well as its ability to escape via a novel non-lytic mechanism known as vomocytosis. We have been exploring whether viral infection affects the interaction between *C. neoformans* and macrophages. Here we show that viral infection enhances cryptococcal vomocytosis without altering phagocytosis or intracellular proliferation of the fungus. This effect occurs with distinct, unrelated human viral pathogens and is recapitulated when macrophages are stimulated with the anti-viral cytokine interferon alpha (IFNα). Importantly, the effect is abrogated when type-I interferon signalling is blocked, thus underscoring the importance of type-I interferons in this phenomenon. Our results highlight the importance of incorporating specific context cues while studying host-pathogen interactions. By doing so, we found that acute viral infection may trigger the release of latent cryptococci from intracellular compartments, with significant consequences for disease progression.

**Non-Technical Author Summary:** Infectious diseases are typically studied in the laboratory in isolation, but in real life people often encounter multiple infections simultaneously. Here we investigate how the innate immune response to the fatal fungus *Cryptococcus neoformans* is influenced by viral coinfection. Whilst virally-infected macrophages retain a normal capacity to engulf and kill Cryptococci, they demonstrate a dramatically enhanced propensity to expel them via the process known as non-lytic expulsion or vomocytosis. Activation of vomocytosis is independent of the type of virus encountered, since both HIV and measles (two entirely unrelated viral pathogens) trigger the same effect. Instead it is driven by interferon-α, a generic ‘antiviral’ response, which signals back to the infected macrophage, triggering expulsion of the fungus. We propose that this hitherto unobserved phenomenon represents a ‘reprioritisation’ pathway for innate immune cells, by which they can alter the frequency with which they expel one pathogen (*Cryptococcus*) depending on the level of threat from a secondary viral infection.

## Introduction

Since their discovery in 1957 by Isaacs and Lindenmann (1), the antiviral effects of type I interferons have been well documented (2–4). More recently, their roles in non-viral infections have been investigated (5, 6). Different bacterial stimuli have been shown to elicit type I interferon production, and in turn these so called “antiviral cytokines” play a role in the outcome of bacterial infections (7–9). This stems in part from the complex and sometimes contradictory effects that type I interferons have on host cells, for instance in enhancing inflammatory responses in some infectious settings (6) to preventing hyperinflammation in others (10, 11), and even affecting the priming of immune responses at lymph nodes (12).

To date, little is known about the interplay between type I interferons and fungal infections, despite the fact that many life-threatening fungal infections occur in the context of chronic viral infection. This is particularly true of *Cryptococcus neoformans*, a globally distributed opportunistic pathogen that is responsible for nearly 200,000 deaths per year in human immunodeficiency virus (HIV) infected people, where it causes cryptococcal meningitis (13). Extensive work over many years has demonstrated that a key feature of cryptococcal pathogenesis is the ability of the fungus to survive, proliferate within, and then escape from, host macrophages (14–17). Macrophages are among the first immune cells to encounter the fungus within the human host (18), and thus are very important in the fight against this pathogen. These cells are able to phagocytose and contain the threat, as happens in immunocompetent hosts, but can also by hijacked by Cryptococcal cells and used as a “Trojan horse” to disseminate to distal sites within the body, particularly to the central nervous system (19). Engulfed Cryptococcal cells can escape from host macrophages through lytic or non-lytic mechanisms, the latter being known as vomocytosis or non-lytic extrusion (20, 21). Most studies to date have focused on the interaction of Cryptococcus with healthy host cells, and consequently how this intracellular lifestyle may be impacted by viral coinfection remains unknown.

Here we show that viral infections enhance vomocytosis of Cryptococci from infected macrophages, without affecting phagocytosis or intracellular proliferation rate of the fungus. This effect is lost when signalling through the type I interferon receptor is blocked, and can be recapitulated by addition of exogenous IFNα. Thus, antiviral responses by the host have a hitherto unexpected impact on the release of intracellular pathogens by vomocytosis.

## Materials and Methods

All reagents were purchased from SIGMA unless otherwise stated.

### *Cryptococcus* Strains

Cryptococcal strains were grown in Yeast Peptone Dextrose (YPD) broth (2% glucose, 1% peptone and 1% yeast extract) at 25°C on a rotator (20 rpm). Yeast from overnight cultures were centrifuged at 6500 rpm for 2 minutes and resuspended in PBS at the required concentration. All experiments were carried out using *C. neoformans var. grubii* serotype A strain Kn99α. Wildtype, GFP- (22) or mCherry-expressing (23) derivatives of Kn99α were used, as stated for each figure.

### Virus strains

HIV-1 virus stocks were generated by transfection of human embryonic kidney 293T cells (European Collection of Authenticated Cell Cultures) as previously described (24, 25). The R9HIVΔ*env* virus was derived from clade B HIV-1 strain (NL43) with 500bp deletion in *env*, pseudotyped with vesiculostomatitis virus G envelope. SIV3mac single round virus like particles (VLPs) containing vpx (SIV3vpx) were generated by transfection into 293T cells with pSIV3+ and pMDG plasmids (26, 27). At 48, 72h and 96h viral containing supernatant was harvested, centrifuged at 800 x g for 10 min and filtered through 0.45 um filter then centrifuged on a 20% sucrose cushion at 20,000 x g for 2h at 4°C. Purified virus was then re-suspended in RPMI media and frozen at −80°C. To quantify single round HIV infection, a vial was thawed for each harvest and serial dilutions used to infect CCR5/CD4 and CXCR4/CD4 transfected NP-2 cells. At 72h post infection wells were fixed in ice cold acetone-methanol and infected cells were identified by staining for p24 protein using a 1:1 mixture of the anti-p24 monoclonal antibodies EVA365 and EVA366 (NIBSC, Center for AIDS Reagents, UK). Infected cells were detected by light microscopy to provide a virus titre (focus-forming U/mL). The SIV3vpx particles were quantified after thawing using a reverse transcriptase (RT) assay colorimetric kit (Roche) following the manufacturer’s instructions to provide a RT ng/mL titre.

Recombinant MeV strain IC323 expressing green fluorescent protein (MeV-GFP) was generated as previously reported by Hashimoto *et al*. (28) MeV-GFP represents a virulent field isolate from Japan (Ichinose-B (IC-B) strain) and was isolated from a patient with acute measles in 1984 (29). For the generation of virus stocks, Vero (ATCC CCL-81) cells overexpressing human SLAMF1 receptor (vero-hSLAM cells) were grown in T75 tissue culture flasks to approximately 80% confluency in DMEM supplemented with 0.4 mg/mL G418. Flasks were infected with MeV-GFP at an MOI of 0.01:1 in 5 mL media for 1 hour at 37°C. After 1h a further 10 mL of DMEM supplemented with 10% FBS was added and infection allowed to continue for 48 h. At harvest the flasks were frozen to −80°C. After thawing, the collected supernatants were centrifuged at 2500 rpm for 10 min at 4°C to pellet cell debris. Aliquoted virus in supernatant was then frozen to −80°C. MeV-GFP viruses were then titred using the TCID-50 method. Vero-hSLAM cells were seeded into flat-bottomed 96 well plates and infected with serial dilutions of thawed MeV-GFP in triplicate. After 72 h, wells were scored for positive or negative infection under UV illumination on a Nikon TE-2000 microscope.

### Ethics Statement

All work with human tissue was approved by the University of Birmingham Ethics Committee under reference ERN_10-0660. Samples were collected specifically for this work and were not stored beyond the duration of the experiments described herein. All donors provided written consent prior to donation.

### Human macrophage isolation and culture

20-40 mL of blood were drawn from healthy donors by venepuncture. 6 mL of whole blood were carefully layered on top of a double layer of Percoll (densities of 1.079 and 1.098 g/mL). Samples were centrifuged in a swing bucket rotor at 150g for 8 minutes, followed by 10 minutes at 1200g, with acceleration and break set to zero. The resulting white disc of peripheral blood mononuclear cells (PBMC) was transferred to a clean vial and incubated with red blood cell lysis buffer at a ratio of 1:3 for 3 minutes, with gentle mixing throughout to prevent clot formation. Cells were then washed with ice cold PBS twice, with centrifugation at 400g for 6 minutes in between each wash, and counted with a haemocytometer. 1×10^6^ PBMC were seeded onto 48-well plates in RPMI-1640 media containing 1% penicillin/streptomycin, 5% heat-inactivated AB human serum and 20 ng/mL M-CSF (Invitrogen). Cells were washed with PBS and resuspended in fresh media on days 3 and 6 of differentiation. Macrophages were ready to use on day 7. A yield of 1×10^5^ macrophages per well was estimated.

### *Cryptococcus* infection

Fungi were opsonised with 10% human AB serum or 18B7 antibody (a kind gift from Arturo Casadevall) for 1 hour and then added to macrophages at a multiplicity of infection of 10:1. Infection was carried out in serum free-media, at 37°C with 5% CO_2_. After 2 hours, cells were washed 3 times with PBS to remove any extracellular fungi and fresh serum free-media was added.

### Drug treatments

Exogenous compounds were added to macrophages at two stages; when infecting with *Cryptococcus* and again when replenishing with fresh media after removing extracellular fungi.

Compounds tested include interferon alpha (IFNα) at concentrations ranging from 5 to 100 pg/mL (Bio-Techne), polyinosinic-polycytidilic acid (polyIC) at 3 and 30 ng/mL (Invivogen), type-I interferon receptor inhibitor (IFNARinh) at 2.5 μg/mL (pbl assay science).

### Co-infection assay

Human monocyte-derived macrophages were infected with either attenuated human immunodeficiency virus (HIV) or MeV-GFP as follows:

For attenuated HIV co-infections, 24h before cryptococcal infection, human monocyte-derived macrophages were infected either with R9HIVΔ*env* at a MOI of 10:1, SIV3vpx at 3 ng/mL or both in serum free RPMI. At 24 h post infection duplicate wells were fixed in ice cold acetone-methanol and infected cells were identified by staining for p24 protein as described above. Experimental wells were infected with antibody opsonised-*Cryptococcus* Kn99α-GFP for 2 hours, washed to remove extracellular fungal cells, and replenished with fresh serum free-media.

Alternatively, macrophages were infected with MeV-GFP at an MOI of 5:1 in serum free-media and kept at 37°C with 5% CO_2_. After 24 hours, cells were washed with PBS and fresh media, supplemented with 5% heat-inactivated human AB serum, was added. After 3 days, cells were co-infected with serum opsonised*-Cryptococcus* Kn99α-mCherry for 2 hours, washed to remove extracellular fungal cells, and replenished with fresh serum free-media.

### Live imaging

Infected samples were kept at 37°C with 5% CO_2_ in the imaging chamber of a Ti-E Nikon Epifluorescence microscope. Images were taken every 5 minutes over an 18-hour period and compiled into a single movie file using NIS Elements software. Movies were blinded by a third party before manual scoring for phagocytosis of *Cryptococcus*, virus infection rates, vomocytosis events, intracellular proliferation rates and macrophage integrity.

### Growth curve assay

A 10-fold diluted cryptococcal overnight culture was inoculated into YPD broth in a 48-well plate (final dilution in well: 1000-fold), in the presence or absence of type-I interferons. The plate was sealed with a breathable membrane and incubated at 37°C within a fully automated plate reader (FLUOStar, BMG Omega). Optical density readings at 600 nm were taken every 30 minutes over a 24 hour-period, with orbital shaking in between readings.

### Data analysis

Statistical analysis was performed using GraphPad Prism 6. Categorical data of phagocytosis or vomocytosis occurrence in the different conditions was assessed using Chi^2^ test and Fisher’s exact test. If data was normally distributed as assessed by Shapiro-Wilk test, then it was compared using Student’s t test. Figures show percentage of *cryptococcus*-infected macrophages experiencing at least one vomocytosis event within each experiment. For intracellular proliferation rates, data was analysed using Mann-Whitney test. Growth curves were fitted to sigmoidal curves and the parameters were compared using Kruskal-Wallis test. All data shown corresponds to at least three independent experiments.

Raw data (collated manual counts for multiple timelapse movies) are provided as supplemental material for each figure. Original timelapse movies, upon which manual scoring was performed, are freely available upon request from the authors.

## Results

Given the relevance of cryptococcosis to HIV+ patients (13), we set out to test whether HIV infection had an effect on vomocytosis of *C. neoformans*. Human monocyte-derived macrophages were infected with HIV-1 capable of a single-round of infection and subsequently with *C. neoformans* and then used for time-lapse imaging over 18 hours. Subsequent scoring showed that virally infected cells had a significantly higher occurrence of cryptococcal vomocytosis (Figure 1A), whilst fungal uptake and intracellular proliferation were unaltered (Figure 1C, 1E).

**Fig 1.**
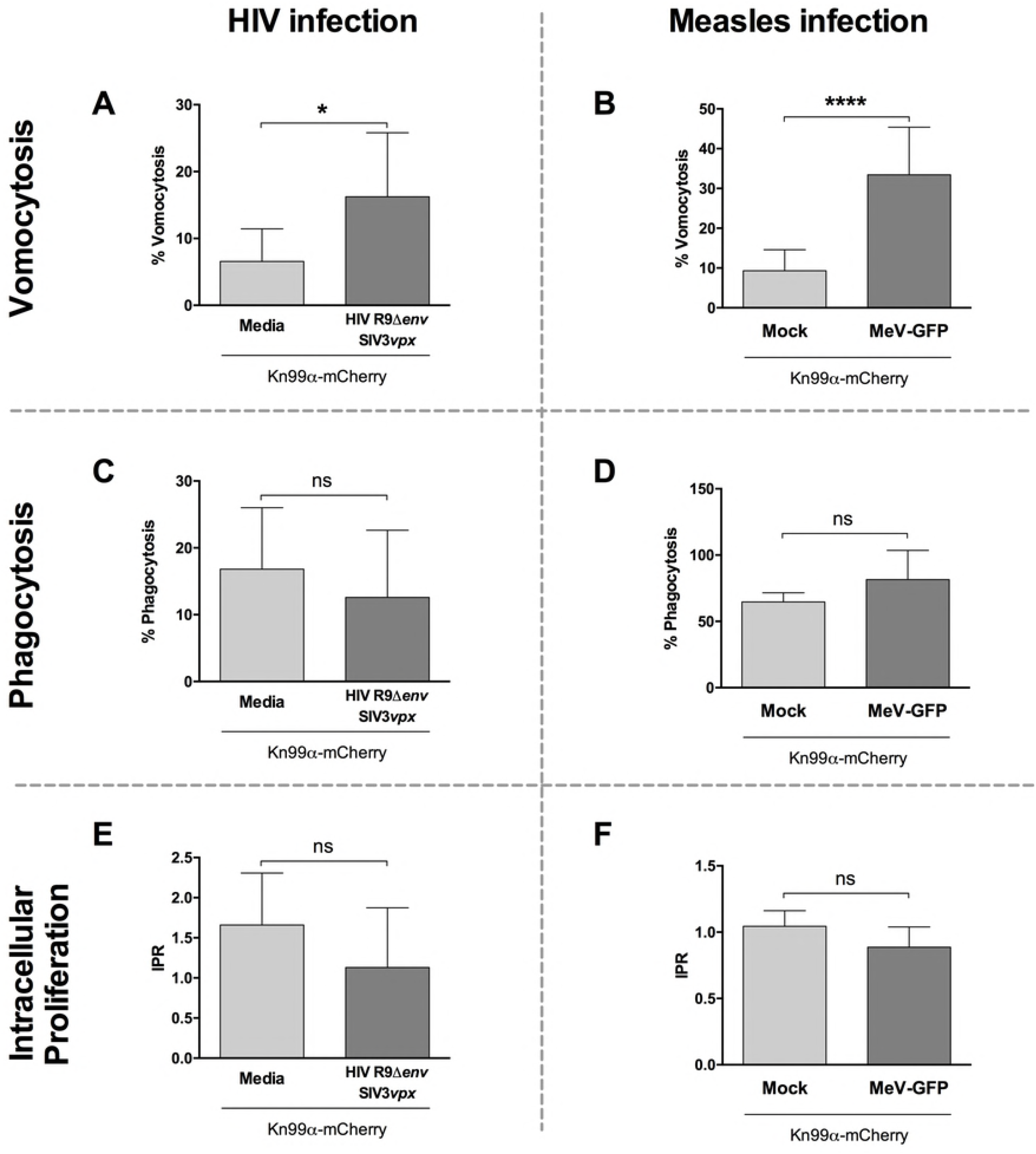
Viral infection enhances vomocytosis of *C. neoformans*. Human monocyte-derived macrophages were infected with HIV (left) or measles virus (right) and subsequently infected with *C. neoformans*. Time-lapse microscopy videos were manually scored for vomocytosis (top), uptake (middle) and intracellular proliferation rate of *C. neoformans* (bottom). **A-B** Graphs show percentage of *cryptococcus-infected* macrophages which have experienced at least one vomocytosis event. **C-D** Percentage of *crytptococus*-infected macrophages. **E-F** Intracellular proliferation rate of *C. neoformans* over 18 hours. In all cases, data corresponds to at least 3 independent experiments. Categorical vomocytosis and phagocytosis data was analysed by Chi^2^ test followed by Fisher’s exact test. * p < 0.05; **** p < 0.0001. IPR data was analysed using Mann-Whitney test.

The experimental HIV system we used here includes co-transduction with SIV3vpx VLPs in order to counteract the antiviral effect of SAMHD1 and ensure maximal HIV infection of the macrophages (26, 30) (Figure S1A). Interestingly, we noted that the addition of SIV3vpx or R9HIVΔ*env* alone also increased vomocytosis (Figure S1B). Since neither condition results in widespread viral infection of host cells, this suggested that the enhancement of vomocytosis occurs at the level of viral detection, rather than being a consequence of active HIV infection.

To explore this further, we tested whether vomocytosis was altered in macrophages infected with an unrelated macrophage-tropic virus (31); measles (MeV, Figure 1B). The measles strain used represents a virulent field isolate from Japan. Once again, infection with the virus resulted in significantly enhanced vomocytosis of *Cryptococcus*. Neither HIV nor measles infection affected uptake of *Cryptococcus* nor the intracellular proliferation rate (IPR) of the fungus (Figure 1C-F), suggesting that the viral effect acts specifically at the level of vomocytosis, rather than fungal pathogenicity *per se*, and that it is independent of the type of virus.

To test whether active viral infection was required for enhanced vomocytosis, we mimicked the effect of viral exposure by stimulating macrophages with polyinosinic-polycytidilic acid (polyIC). PolyIC is a double-stranded RNA synthetic analogue, which is known to trigger antiviral responses by binding to TLR3 (32). Human monocyte-derived macrophages were stimulated with polyIC and infected with *C. neoformans* simultaneously. Infected cells were imaged over 18 hours and scored for vomocytosis (Figure 2A). As with HIV or MeV infection, polyIC stimulation enhanced vomocytosis of *Cryptococcus*. Thus, it is likely that the antiviral reaction of the host macrophage, rather than an aspect of viral pathogenesis, is the trigger for enhanced vomocytosis from infected host cells.

**Fig 2.**
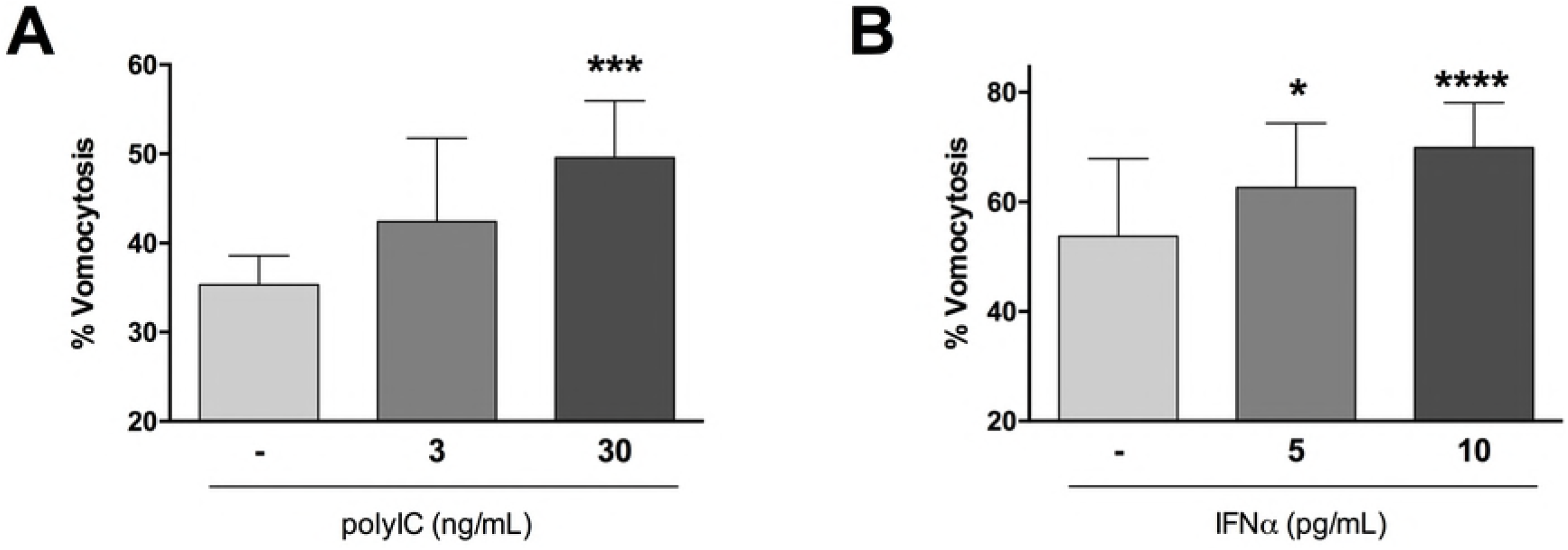
Antiviral response increases vomocytosis. Human monocyte-derived macrophages were stimulated with different doses of polyIC (**A**) or IFNα (**B**), and infected with *C. neoformans*. Graphs show Mean + SD of percentage of *cryptococcus*-infected macrophages which have experienced at least one vomocytosis event. Chi^2^ test followed by Fisher’s exact test performed on raw vomocytosis counts. Data corresponds to at least three independent experiments.

The hallmark of the cellular anti-viral response is the induction of type-I interferons. Among these, the best studied are IFNα and IFNβ. During HIV infection specifically, the induction of IFNα is the most relevant (33). We therefore tested whether the impact of viral infection on vomocytosis could be recapitulated by exposure to interferon-α (IFNα). Stimulation of human monocyte-derived macrophages with 10 pg/mL IFNα (a level that closely matches that seen in HIV-infected patients (33)) resulted in significantly enhanced vomocytosis of *Cryptococcus*(Figure 2B) without altering cryptococcal growth, uptake or IPR (Figure S2). Interestingly, we noticed that higher doses of IFNα suppressed this effect, suggesting that the impact of interferons on vomocytosis can be rapidly saturated.

To confirm that type-I interferons were behind the increase in vomocytosis observed, we performed the viral infection experiments in the presence of a type-I interferon receptor (IFNAR) inhibitor (Figure 3). The addition of IFNAR inhibitor blocked the enhancement of vomocytosis otherwise elicited by viral infection in both HIV- and Measles-infection settings, confirming that type-I interferon signalling is necessary for this effect. Interestingly, this effect was particularly prominent on virally infected cells rather than neighbouring cells which were not infected (Non-MeV; Figure 3B), suggesting that the impact of IFNα signalling on vomocytosis is highly localised and specific to the autocrine responses occurring within infected cells, rather than endocrine responses mediated through cytokines.

**Fig 3.**
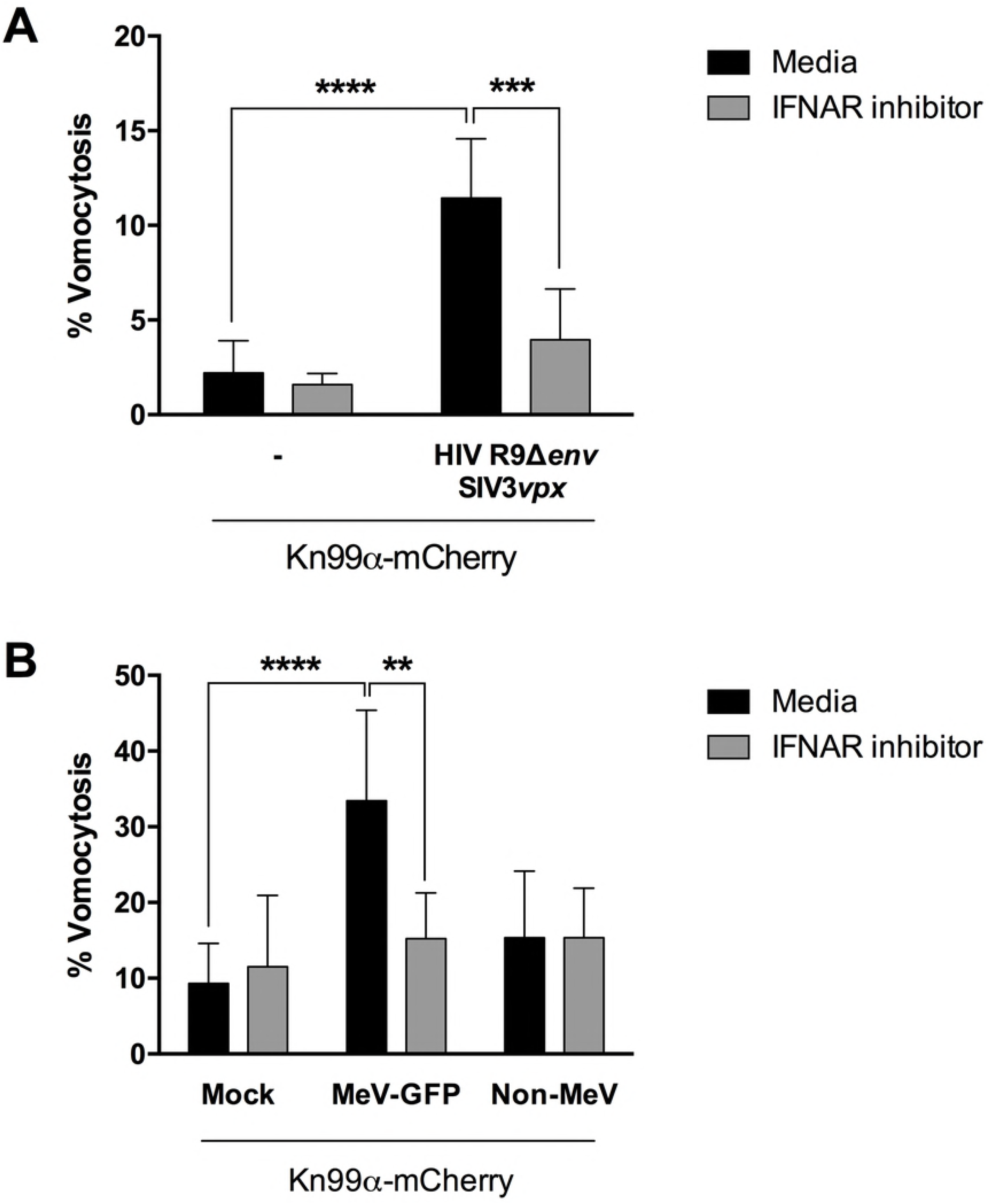
Type-I interferon signalling is necessary to enhance vomocytosis. Human monocyte-derived macrophages were infected with HIV (**A**) or GFP-expressing measles virus (MeV-GFP, **B**) and subsequently with mCherry-expressing *C. neoformans*(Kn99α-mCherry), in the presence or absence of an IFNAR blocking antibody. GFP negative cells, which did not have an active Measles infection, were termed “Non-MeV”. Graph shows Mean + SD of percentage of *Cryptococcus*-infected macrophages which have experienced at least one vomocytosis event. Fisher’s exact test performed on raw vomocytosis counts. Data corresponds to two and three biological repeats, respectively.

## Discussion

In this study we set out to explore the consequences, if any, of viral infection on Cryptococcal infection, focusing on the non-lytic escape mechanism known as vomocytosis. Infection with either HIV or measles virus led to an enhancement in vomocytosis of *C. neoformans*, without affecting uptake or intracellular proliferation of the fungus (Figure 1), an effect that could be recapitulated by stimulation with IFNα and abrogated when signalling from type-I interferon receptor was blocked (Figures 2 and 3). Thus, viral coinfection stimulates expulsion of intracellular fungi via Type I interferon signalling.

The effect was seen using two distinct viral pathogens which differ, among other parameters, in the magnitude of anti-viral response they elicit in human macrophages. Relative to other viral infections, HIV is very good at avoiding the induction of type-I interferons (24, 25). Nonetheless, the low levels of type-I interferons induced by HIV, potentially enhanced by the co-infection with *Cryptococcus*, are sufficient to have a significant effect on vomocytosis. Infection with measles virus has been reported to induce limited production of type-I interferons in macaque models, albeit with potent induction of interferon-stimulated genes (34, 35). To date, there is no direct correlation between measles infection and cryptococcosis. However, given that both pathogens have a distinct respiratory phase it is possible that they interact within this shared niche, potentially through low doses of antiviral signalling.

Why might antiviral signalling induce vomocytosis? One possibility is that vomocytosis serves to “reset” phagocytes that have been unable to kill their prey, thus allowing them to serve a useful purpose in phagocytosing other pathogens rather than remaining “unavailable”. In that context, a potent inflammatory signal such as IFNα may serve to accelerate this process during localised infection, returning macrophages to functionality faster than would otherwise occur. The consequences of vomocytosis on disease progression, however, are likely to be highly context dependent; in some settings, this may enable a more robust immune response, but in others it may serve to inadvertently disseminate the fungus to distal sites.

This is supported by previous reports showing variable outcomes of interferon signalling on cryptococcal infection in mice. Sato *et al*. (36) showed that IFNARKO mice have lower fungal burden than WT mice and consequently argue that type-I interferon signalling is detrimental for the host during cryptococcal infection. Supporting this notion but using the sister species *C. gatti*, Oliveira *et al* (37) show that infection with influenza virus worsens the prognosis of subsequent fungal infection. On the other hand, Sionov *et al* (38) showed that stimulation with IFNα or with the double-stranded RNA analogue pICLC protected the host from infection by either *C. neoformans* or *C. gatti* infection. This effect was time-dependent, with the protective effect of pICLC treatment only occurring if administered during the first 72 hpi before the fungus reaches the brain. A tempting model, therefore, is that stimulating vomocytosis via antiviral signalling early in infection (when the fungus remains in the lung) helps prevent dissemination, whilst triggering vomocytosis later on may actually enhance fungal spread and accelerate disease progression.

Taken together, our findings therefore suggest that the antiviral response, and IFNα in particular, induce the expulsion of intracellular cryptococci and that this effect could be advantageous or detrimental to the host, depending on the localization of the infected phagocyte and timing of the event.

## Acknowledgments

PIS, LMTS and RCM are supported by funding from the European Research Council under ‘ the European Union’s Seventh Framework Programme (FP/2007-2013)/ERC Grant Agreement No. 614562. RCM holds a Wolfson Royal Society Research Merit Award and PIS is supported by a Darwin Trust scholarship. MN is supported by a Wellcome Trust Investigator Award (207511/Z/17/Z) and the NIHR Biomedical Research Centre at University College London Hospitals.

## Supporting information

**Fig S1**

**A.** Human monocyte-derived macrophages were infected with VLPs as indicated. After 24 hours, viral infection was assessed by p24 staining (blue).

**B.** Cells were infected with VLPs as indicated, and subsequently infected with *C. neoformans*.Time-lapse microscopy videos were manually scored for vomocytosis. Graph shows percentage of *cryptococcus*-infected macrophages which have experienced at least one vomocytosis event. Chi^2^ test followed by Fisher’s exact test performed on raw vomocytosis counts from 5 independent experiments.

**Fig S2**

**A.** Cryptoccocal cells were grown in the presence or absence of IFNα over 24 hours. Growth was assessed by optical densitiy readings at 600 nm.

**B-C.** Human monocyte-derived macrophages were infected with *C. neoformans* in the presence of different doses of recombinant IFNα. Time-lapse microscopy videos were manually scored for phagocytosis and intracellular proliferation rate of the fungus (B and C, respectively).

Data corresponds to 3 independent experiments.

